# Senescence associated protein degradation

**DOI:** 10.1101/100388

**Authors:** Gerardo Ferbeyre

## Abstract

Senescent cells accumulate with age and contribute to pathologies associated to old age. The senescent program can be induced by pro-cancer stimuli or is developmentally controlled. In cells forced to senesce by expression of oncogenes or short telomeres, aberrant activation of the ERK/MAP kinase signaling pathway leads to selective protein degradation by the ubiquitin proteasome system. The proteins affected by this process control key cellular processes known to be defective in senescent cells. We discuss the evidence supporting a general role for senescence associated protein degradation for organismal aging.

## Introduction

Organisms progressively decay, wear out and die. However, the primary causes triggering these processes have been largely elusive. At the cellular level, cells can die or senesce and the mechanisms of these cellular fates can give insights into organismal aging. Cellular senescence can be greatly accelerated by oncogenic signalling which is different from normal signalling in strength and response to negative modulators. Oncogenic signalling is thus aberrant signalling. Oncogenic *ras*, for example, activates the ERK and AKT kinases leading to the phosphorylation of multiple proteins. Using proteomics analysis, we found that aberrant ERK signalling due to expression of oncogenic *ras* or short telomeres leads to degradation of specific proteins (Deschenes-Simard and others 2013). Most of the degraded proteins contained phosphorylation motifs for proline directed kinases such as ERK1, ERK2, CDKs and GSKs. These kinases are stimulated by oncogenic *ras* or short telomeres and provide a direct link between senescent stimuli and protein degradation. In addition, many unstable proteins in senescent cells were found to contain phosphorylation sites for basophilic kinases such as AKT and S6 kinase or acidophilic kinases such as casein kinases 1 and 2. These kinases could be stimulated as a consequence of signaling by Ras-activated kinases. Alternatively, ERK-dependent production of reactive oxygen species can enhance overall protein phosphorylation due to inhibition of protein phosphatases (Meng and Zhang 2013). The proteins degraded in senescent cells play roles in cell cycle progression, ribosome biogenesis, cell migration, mitochondria and other functions known to be affected in these cells (Deschenes-Simard and others 2014).

The stoichiometry of most phosphorylation events indicates that only a fraction of a particular protein is phosphorylated (Olsen and others 2010). Therefore, during normal cell signaling the coupling between protein phosphorylation and degradation will only impact a fraction of any particular protein pool. However, during aberrant signaling, as in cells bearing constitutively active oncogenes, a larger fraction of some particular proteins will be phosphorylated and subsequently degraded. We call this process Senescence Associated Protein Degradation or **SAPD** (Deschenes-Simard and others 2013) and for some proteins it dominates over their biosynthesis, significantly reducing their overall levels. This mechanism is independent of cell division and could in principle explain cell dysfunctions for both dividing and non-dividing cells exhibiting aberrant signaling.

Oncogenic mutations are not linked to normal aging but senescent cells accumulate with age in many organisms including humans (Childs and others 2015; Tchkonia and others 2013). On the other hand, cells could senesce *in vivo* in response to still unknown factors triggering aberrant signaling and SAPD (Fig 1). The SAPD model of senescence includes: 1) a cause for aberrant signalling, 2) coupling protein modifications induced by aberrant signaling to protein degradation and 3) the cellular consequences of depletion of SAPD target proteins. Here, we review the evidence that links aging and the SAPD model.

**Fig 1.**
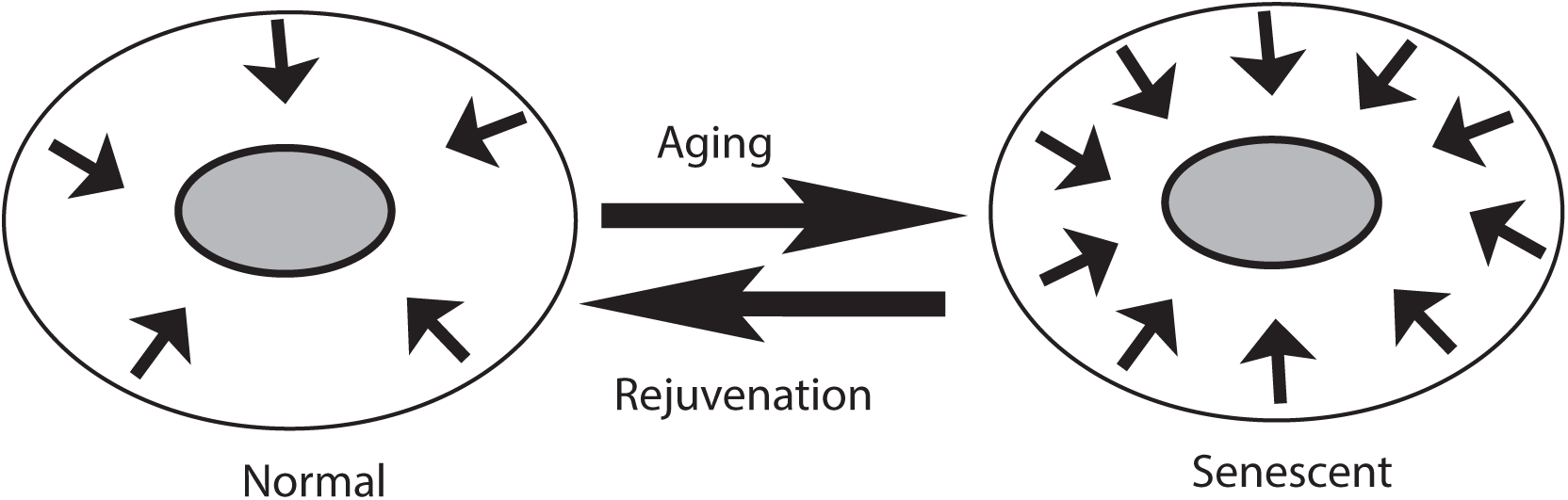
Aberrant signaling underlies cell senescence and aging. Senescent cells in old organisms have a hyperactivation of signaling pathways due to triggers that remain to be discovered. In cell culture, short telomeres, oncogenes and DNA damaging drugs can mimic the process. In animal models mutations or compounds that reduce signaling increase life span.

### 1. Aberrant signaling during aging

Signaling includes a variety of protein modifications including phosphorylation, acetylation, Methylation and sumoylation. These modifications may all contribute to aging but here we focus on protein phosphorylation which is the best understood. There is an extensive body of work on protein acetylation and aging, a pathway controlled by the protein deacetylases of the sirtuin family, which has been reviewed elsewhere (Haigis and Sinclair 2010) and will not be included here.

#### 1.1 Model organisms

Genetic analysis of aging in model organisms has lead to the identification of kinases that shorten life span. In *C. elegans* the insulin/IGF-1 signaling activates the PI3K/AKT cascade leading to phosphorylation and cytoplasmic sequestration of the forkhead transcription factor Daf-16 (Paradis and Ruvkun 1998). This transcription factor is a key mediator of longevity in worms and it regulates the expression of genes that can increase resistance to a variety of cellular stresses (Lee and others 2003). Intriguingly, Daf16 also acts as a transcriptional repressor of many protein kinase genes including those that inactivate its own function (Tazearslan and others 2009). This striking attenuation of many kinase-signaling modules included the PI3 kinase pathway, the TOR pathway and the ERK/MAP kinase pathway (Tazearslan and others 2009). Clearly, protein phosphorylation is linked to aging and its attenuation increase life span (Fig 1).

Consistent with the work in worms, inhibitors of the ERK pathway (Slack and others 2015) or mutations in the PI3K/AKT pathway (Yamamoto and Tatar 2011) also prolonged life span in flies. Moreover, in agreement with the idea for a pro-longevity role for kinase attenuating mechanisms, fibroblasts obtained from long-lived mutant mice, including the Snell dwarf mice and the Growth Hormone Receptor mutant mice, displayed an attenuated ERK activation in response to oxidative stress (Sun and others 2009).

Another protein kinase that has pro-aging functions is casein kinase 1. This kinase is controlled by the proteasome activator REGg. Mice null for REGg accumulate CK1, which then phosphorylates Mdm2, targeting it for proteasome-dependent degradation (Inuzuka and others 2010). As a result, these mice experience an increase in p53 levels and activity and premature aging (Li and others 2013). CK1 is activated by DNA damage and colocalize with p53 in PML bodies where it phosphorylates p53 at threonine 18 an event that prevents its interaction with Mdm2 (Alsheich-Bartok and others 2008). Signs of CK1 activation in Alzheimer disease also suggest a role for this kinase in human aging (Flajolet and others 2007; Hanger and others 2007).

#### 1.2 Human aging

Aberrant phosphorylation of the neuronal cytoskeleton by ERK kinases or decrease in phosphatases is associated to brain aging and human Alzheimer disease (Veeranna and others 2004; Veeranna and others 2011). MEK inhibitors prevent memory deficits in a mouse model of Alzheimer disease (Feld and others 2014). The Raf-1 kinase, which acts upstream ERK1/2 in the MAPK pathway, is also increased in the brain of Alzheimer disease patients (Mei and others 2006), and treatment with Raf kinase inhibitors protects cortical brain cells from β-amyloid toxicity (Echeverria and others 2008). Raf inhibitors also reduced mutant huntingtin toxicity in a cell model of Huntington disease where RNAi-mediated inhibition of multiple members of the RAS-RAF-ERK pathway rescue cells and animal models form this toxicity (Miller and others 2012). GSK3b is another kinase that can phosphorylate the cytoskeletal protein tau in neurons from patients with Alzheimer disease (Noble and others 2005). This kinase increases with aging in the brain and accumulates in the nucleus of senescent fibroblasts (Zmijewski and Jope 2004). The most common genetic mutation leading to Parkinson disease affects the protein kinase LRRK2, and most of its effects can be explained by activation of the ERK pathway. Treatment with MEK inhibitors reversed most of the phenotypic effects of this mutation in neuron cultures (Bravo-San Pedro and others 2013; Carballo-Carbajal and others 2010; Reinhardt and others 2013; White and others 2007). As reported for Alzheimer disease, a reduction in protein phosphatase 2A activity can also underlie an aberrant protein phosphorylation state in Parkinson disease (Wu and others 2012).

According to the SAPD model attenuation of kinase signaling should extend life-span by preventing the conversion of signaling pathways from a state of moderate signaling to a state of aberrant signaling leading to protein inactivation by SAPD (Fig 1). Organisms need to use these signaling modules for cell proliferation and growth and they are in constant danger of passing the threshold for aberrant signaling, SAPD and senescence. A trade must be reached and longer life span can be achieved by attenuating signaling pathways. However, there is a fitness cost associated to lower activity that depends on the environmental pressures and the competition within species to pass genes to the next generation.

#### 1.3 Mechanism of aberrant kinase activation during aging

The reason for aberrant kinase activation in aging is not known. It is unlikely that mutations such as those found in oncogenic *ras* in tumors mediate aberrant kinase stimulation during aging. One possible explanation is the emergence of aberrant signalling by epigenetic mechanisms encoded in the dynamics of phosphorylation cascades. Protein kinases can inherently sustain a persistent activation if they autophosphorylate (Lisman 1985; Song 2013). Calpain, a protease linked to Alzheimer disease, can generate a hyperactive form of GSK-3β after proteolytic cleavage of its C-terminus (Jin and others 2015). Kinase signaling pathways can locked into permanent activated states by triggering positive feedback mechanisms (Lisman and Fallon 1999). For example, ERK can activate PKC via PLA2 and in turn, PKC can activate ERK via RAF-MEK (Lisman and Fallon 1999; Xiong and Ferrell 2003). Positive feedback mechanisms are ideal to mediate stable cellular states because the information is not lost when one component of the loop is degraded. Newly synthesized components are readily integrated into the mechanism by the other members of the loop. Phosphorylation-dependent feed back loops are modulated by protein phosphatases that set a threshold for their activation. Inactivation of phosphatases facilitates the establishment of kinase-based memory circuits (Sweatt 2001) but also impair the process of erasing memories (Silva and Josselyn 2002).

Repeated stimulation of cells with cytokines causes persistent ERK activation that depends on a positive feedback loop involving Sprouty 2. Sprouty 2 activates the tyrosine kinase Fyn, which is capable of activating ERK (Liu and others 2010). This Sprouty-dependent activation of ERK was associated to accumulation of activated ERK in endosomes. Further identification and characterization of positive feedback modules within signalling pathways will help to discover those driving aberrant kinase activation during aging and provide targets for pharmacological intervention. Senescent cells secrete a variety of pro-inflammatory modulators that can reinforce the senescence state in a cell autonomous manner and propagate the senescent phenotype to neighbouring cells (Acosta and others 2013; Acosta and others 2008a; Acosta and others 2008b). These inflammatory cytokines act in part by inducing the generation of reactive oxygen species and stimulating signaling pathways including the ERK pathway (Acosta and others 2008a).

It has been argued that the hyperfunctional signaling associated to senescence is the continuation of developmental programs that are not switched off (Blagosklonny 2013). These programmes are optimized for embryonic development and growth but their persistence in the adult somehow leads to aging. In fact, cellular senescence is used as a mechanism to eliminate and replace certain embryonic structures during development (Munoz-Espin and others 2013; Storer and others 2013) and to halt the expansion of potentially malignant cells (Serrano and others 1997). Therefore, although genes do control development, senescence and aging there is no evidence that they were selected for their pro-aging functions. Aging, in this view, is a side effect of developmental programs (“a shadow of actual programs’) (Blagosklonny 2013) and/or the price of tumor suppression (Ferbeyre and Lowe 2002).

### 2-Coupling protein modifications to protein degradation

In many situations, phosphorylation is coupled to protein degradation and in general phosphoproteins have a higher turnover. This coupling reset signalling modules and allows cells to respond to different stimuli or repeated stimulation (Cambridge and others 2011). Various kinase motifs increase protein turnover. For example proteins having ERK-kinase motifs are less stable than proteins having AKT motifs (Cambridge and others 2011). Also, multiple ubiquitin conjugating enzymes recognize only phosphorylated proteins to which they attach ubiquitin chains for proteasome-dependent degradation (Harper 2002). Phosphorylation can also inactivate signaling pathways by controlling protein localization (Paradis and Ruvkun 1998) or altering protein conformation (Atadja and others 1994).

The degradation of phosphorylated proteins by the proteasome depends on E3 ubiquitin ligases that recognize phosphorylated proteins at specific motifs known as phosphodegrons. The ability of SAPD to deplete key proteins thus depends on the presence of these phosphodegrons in target proteins, E3 ligases that recognizes them and an aberrant signaling that changes the normal stoichiometry of phosphorylation labeling most of a specific protein pool for degradation. E3 ligases of the SCF family possess F-box proteins that bind directly to phosphorylated proteins. Several E3 ligases have been linked to cellular senescence. Smurf2, an E3 ligase in the TGFβ pathway, is activated by short telomeres and mediates cellular senescence (Zhang and Cohen 2004). Smurf2 ubiquitinates Id1 and Id2, two repressors of p16INK4a-expression, explaining the widespread upregulation of p16INK4a in senescent cells (Kong and others 2011). Smurf2 also targets the polycomb protein and epigenetic regulator EZH2 for degradation (Yu and others 2013) a function that should also contribute to p16INK4a upregulation via erasing repressive modifications on the p16INK4a promoter chromatin (Bracken and others 2007).

In nematodes, the E3 ubiquitin ligase RLE-1 reduces longevity by catalyzing the degradation of DAF16, a forkhead family transcription factor (Li and others 2007). The deubiquitylase MATH-33/USP7 reverses the action of RLE-1 and promotes longevity (Heimbucher and others 2015). Another E3 ligase component that promotes aging in nematodes is elongin c. RNAi-dependent downregulation of this gene increases life span by increasing the levels of its target HIF-1, a transcription factor with anti-aging activity (Hwang and others 2015). There are many E3 ubiquitin ligases encoded in the mammalian genome and for the most part they remain poorly characterized. We anticipate that many more E3 ubiquitin ligases and phosphodegrons will be implicated in the regulation of senescence and aging.

### 3-Consequences of senescence-associated protein degradation

Studying protein turnover in several mouse models of extended longevity provided evidence linking protein degradation to aging. Longevity correlated with a reduction in protein turnover but not with a reduction in cell proliferation rates as predicted by the telomere hypothesis. The mechanisms underlying a reduction in protein turnover in long living models remain to be investigated but the authors suggested that they are likely the result in a reduction of protein damage or misfolding (Thompson and others 2016). Their data is not consistent with increased proteolytic editing of damaged proteins as suggested by theories implicating increased autophagy in longevity or decreased proteasome protein degradation in aging.

Among conditions associated to aging protein degradation has been well studied in muscle atrophy (Altun and others 2010). Aged muscle cells contain higher levels of the ubiquitin ligase CHIP and MURF1 (Altun and others 2010), which promote the ubiquitination and degradation of misfolded proteins. In situations leading to muscle atrophy, such as denervation and cachexia, a common transcriptional program is activated to induce the so called atrogenes (Zheng and others 2010). Several components of the ubiquitin-proteasome system including the E3 ligases MURF-1 and atrogin are induced by activation of FOXO transcription factors (Zheng and others 2010). Another important component of the protein degradation machinery during muscle atrophy is the p97/VCP/CDC48 ATPase, which binds multiple E3 ligases and catalyzes ATP-driven disassembly of protein complexes. This ATPase is thus important for extraction of ubiquitinated proteins from protein complexes in ER associated proteins, mitochondrial complexes, myofibrils and chromatin (Piccirillo and Goldberg 2012). Inhibition of p97 reduces muscle atrophy after fasting or denervation (Zheng and others 2010) and its levels increase during aging related sarcopenia (Altun and others 2010). The trigger of muscle atrophy during aging is unknown. However, an intriguing increase in phospho-ERK was observed in resting old muscles in comparison to young muscles (Williamson and others 2003) and the authors suggested that old muscle are under constant stress signaling due to inflammatory cytokines such as TNF, which has been reported to increase with aging in muscles (Kirwan and others 2001).

A transgenic mouse reporter for the ubiquitin proteasome system revealed no significant alterations during aging (Cook and others 2009). Intriguingly, in long living daf2 mutants of C. elegans, translation and proteasome components were reduced (Stout and others 2013). Reduced protein degradation via a decrease in proteasome components allowed normal protein content in these mutants (Stout and others 2013). These results are consistent with the SAPD model of aging. Finally, an increase in phosphorylation-dependent protein degradation may divert the proteasome from its function in the degradation of damaged proteins (Hernebring and others 2006) further compromising the physiology of cells with aberrant signaling.

## Concluding remarks

Aging is obviously associated with deterioration and loss of functions. Paradoxically it can be triggered at the cellular level by hyperfunctional pathways. We propose that the link between hyperfunction or aberrant signaling and the loss of function associated to aging is the degradation of aberrantly modified proteins. The SAPD model thus provides a molecular mechanism to link hyperfunction to the multiple defects associated to senescent cells. The SAPD model of aging can be tested in model organisms where longevity could be linked to downregulation of key protein kinases and/or E3 ubiquitin ligases mediating the process of phosphorylation-dependent protein degradation. Small molecules modulating this pathway could be used for anti-aging strategies.

## Acknowledgements

I thank members of my laboratory for useful comments. The work on senescence is supported by CIHR, FRQS and the Canadian Cancer Society.

